# Comparative analysis of RNA enrichment methods for preparation of *Cryptococcus neoformans* RNA sequencing libraries

**DOI:** 10.1101/2021.03.01.433483

**Authors:** Calla L. Telzrow, Paul J. Zwack, Shannon Esher Righi, Fred S. Dietrich, Cliburn Chan, Kouros Owzar, J. Andrew Alspaugh, Joshua A. Granek

## Abstract

Ribosomal RNA (rRNA) is the major RNA constituent of cells, therefore most RNA sequencing (RNA-Seq) experiments involve removal of rRNA. This process, called RNA enrichment, is done primarily to reduce cost: without rRNA removal, deeper sequencing would need to be performed to balance the sequencing reads wasted on rRNA. The ideal RNA enrichment method would remove all rRNA without affecting other RNA in the sample. We have tested the performance of three RNA enrichment methods on RNA isolated from *Cryptococcus neoformans*, a fungal pathogen of humans. We show that the RNase H depletion method unambiguously outperforms the commonly used Poly(A) isolation method: the RNase H method more efficiently depletes rRNA while more accurately recapitulating the expression levels of other RNA observed in an unenriched “gold standard”. The RNase H depletion method is also superior to the Ribo-Zero depletion method as measured by rRNA depletion efficiency and recapitulation of protein-coding gene expression levels, while the Ribo-Zero depletion method performs moderately better in preserving non-coding RNA (ncRNA). Finally, we have leveraged this dataset to identify novel long non-coding RNA (lncRNA) genes and to accurately map the *C. neoformans* mitochondrial rRNA genes.

**ARTICLE SUMMARY:** We compare the efficacy of three different RNA enrichment methods for RNA-Seq in *Cryptococcus neoformans*: RNase H depletion, Ribo-Zero depletion, and Poly(A) isolation. We show that the RNase H depletion method, which is evaluated in *C. neoformans* samples for the first time here, is highly efficient and specific in removing rRNA. Additionally, using data generated through these analyses, we identify novel long non-coding RNA genes in *C. neoformans*. We conclude that RNase H depletion is an effective and reliable method for preparation of *C. neoformans* RNA-Seq libraries.

## INTRODUCTION

RNA sequencing (RNA-Seq) is a powerful tool for quantifying gene expression in diverse organisms. Despite the rapid and continual decrease in sequencing costs, the expense of sequencing is often the limiting factor in designing RNA-Seq experiments. Due to this cost constraint, enrichment of the RNA classes of interest, hereafter referred to as “RNA enrichment”, is an important step in library preparation for most RNA-Seq experiments. Ribosomal RNA (rRNA) is the most abundant RNA in any cell, generally constituting more than 90% of the total RNA (Giannoukos *et al*. 2012). Despite its abundance, it is rarely of interest in RNA-Seq experiments, therefore, 90% or more of the data is useless when generated without an RNA enrichment step. RNA enrichment aims to reduce the content of RNA in the library, eliminating sequencing capacity wasted on uninformative data and reducing the cost of data storage and analysis, thus decreasing the overall cost of the experiment.

There are many different methods for RNA enrichment and many products available based on these different methods. When selecting an RNA enrichment method and product there are two key considerations: 1) the fraction of rRNA removed; 2) the side effects on other RNAs in the sample. RNA enrichment methods either specifically target the RNA of interest, most commonly mRNA, for isolation or specifically target rRNA for removal (Zhao *et al*. 2014). The most common mRNA isolation method, poly(A) isolation, uses an oligo(dT) affinity matrix. Raw RNA is hybridized to the matrix, which preferentially binds the 3’ polyadenylation sequence of mRNA. By enriching polyadenylated mRNA, rRNA, which lacks 3’ polyadenylation, is depleted de facto. Although mRNA isolation methods are typically efficient in eliminating rRNA, they fail to capture any RNA molecules lacking polyadenylation, such as non-coding RNA (ncRNA). They are also only applicable to eukaryotes, since mRNA in prokaryotes is generally not polyadenylated. Most rRNA removal methods involve hybridization of sequence-specific probes to rRNA. These probes target the rRNA for depletion. In the Ribo-Zero depletion method, the probes are synthesized with a molecular tag, which is used to bind the probe-rRNA complex to beads, allowing the complexed rRNA to be removed from solution (Zhao *et al*. 2014). In the ribonuclease H (RNase H) depletion method, sequence-specific DNA probes hybridized to rRNA target the rRNA for enzymatic degradation by RNase H, which specifically degrades RNA from RNA-DNA complexes (Morlan *et al*. 2012). The duplex-specific nuclease (DSN) method indiscriminately depletes high abundance sequences by denaturing and reannealing the prepared RNA-Seq library, then treating with a duplex-specific nuclease to degrade all double-stranded DNA. Under the conditions used for reannealing, high abundance sequences are much more likely to find a complementary sequence, so high abundance sequences, including but not limited to rRNA, are preferentially removed from the pool (Yi *et al*. 2011).

The poly(A) isolation and DSN methods are attractive because they are broadly applicable without any organism-specific adaptation: the poly(A) method works in all eukaryotes and the DSN method should work in any organism. However, the rRNA removal methods (Ribo-Zero and RNase H) are more targeted, and are therefore expected to have fewer side-effects on biologically-important RNA molecules, such as protein-coding RNA and ncRNA. The downside inherent in the targeted nature of the rRNA depletion methods is that the sequence-specific probes must be designed for the organism under experimentation or a close relative for maximal efficacy. Because rRNA is the most highly conserved sequence across the tree of life (Isenbarger *et al*. 2008), probes designed for an evolutionarily distant species will often work, but efficiency of rRNA depletion decreases with evolutionary distance. For all rRNA depletion methods, the performance of these rRNA removal methods can vary by organism, so it is important to assess them on the organism of interest.

The budding yeast *Cryptococcus neoformans* is a human fungal pathogen that infects more than 200,000 people annually and causes excessive mortality among immunocompromised patient populations, such as those with HIV/AIDS and those receiving immunosuppressive cancer therapies (Rajasingham *et al*. 2017). Research on *C. neoformans* helps us better understand this pathogen, contributes to the development of treatments for *C. neoformans* infections, and advances our understanding of fungal pathogens in general. RNA-Seq has been used extensively in *C. neoformans* studies to elucidate regulatory networks of protein-coding genes (Chang *et al*. 2014; Chen *et al*. 2014; Janbon *et al*. 2014; Gish *et al*. 2016; Chow *et al*. 2017; Brown *et al*. 2018, 2020; Yu *et al*. 2020). While there is intense interest in the role of ncRNA in higher eukaryotes such as humans, relatively little work has explored the implications of ncRNA in fungi, with focus largely on model fungi such as *Saccharomyces cerevisiae* (Bird *et al*. 2006; Bumgarner *et al*. 2009, 2011; Gelfand *et al*. 2011; Parker *et al*. 2018) and *Schizosaccharomyces pombe* (Ding *et al*. 2012; Atkinson *et al*. 2018). However, multiple recent studies have demonstrated the importance of ncRNA in *Cryptococcus* biology and virulence, including microRNA (miRNA) (Jiang *et al*. 2012; Liu *et al*. 2020), small interfering RNA (siRNA) (Wang *et al*. 2010; Janbon *et al*. 2010; Liu *et al*. 2020), and long non-coding RNA (lncRNA) (Fan *et al*. 2005; Chacko *et al*. 2015; Liu *et al*. 2020).

In planning and analyzing RNA-Seq experiments in *C. neoformans*, it is essential to understand the side-effect profile of the RNA enrichment method used. RNA enrichment methods that alter levels of RNA of interest may give misleading or incorrect results; this is a special concern for analysis of ncRNA. Here, we assess three different enrichment methods for RNA-Seq applications in *C. neoformans*: RNase H depletion, Ribo-Zero depletion, and Poly(A) isolation. The Ribo-Zero depletion (“Ribo-Zero Kit Species Compatibility Tables”; Trevijano-Contador *et al*. 2018; Liu *et al*. 2020) and Poly(A) isolation (Bloom *et al*. 2019; Brown *et al*. 2020) methods have been used previously in *C. neoformans*, while the RNase H method has not. However, none of these methods have been evaluated in *C. neoformans* in comparison to each other, much less to an unenriched “gold standard.” By performing this controlled experiment, we have been able to quantify the efficiency of rRNA depletion and determined the side-effects of each depletion method on non-rRNA genes.

We find that the RNase H depletion method is more efficient than the Ribo-Zero depletion and the Poly(A) isolation methods in removing rRNA. Additionally, we report that the RNase H depletion method is highly specific; it performs better than the other two methods in preserving protein-coding RNA, and nearly as well as the Ribo-Zero depletion method in preserving ncRNA. Because the RNase H depletion and Ribo-Zero depletion methods both preserve ncRNA, we utilized the data generated from these methods to identify novel *C. neoformans* lncRNA. Collectively, this work demonstrates that RNase H depletion is an effective RNA enrichment method for use in preparation of *C. neoformans* RNA-Seq libraries, further emphasizes the role of RNA enrichment in design of economical RNA-Seq experiments, and highlights the importance of knowing the side-effect profile when choosing an RNA enrichment method.

## MATERIALS AND METHODS

### Strains, media, and growth conditions

The *C. neoformans* var. *grubii* H99 (*MAT*α) wild-type strain was used for all experiments. This strain was maintained on yeast extract-peptone-dextrose (YPD) medium (1% yeast extract, 2% peptone, 2% dextrose, and 2% agar for solid medium).

### RNA-Seq library preparation

Three biological replicate samples (A, B, and C) were used for all analyses. Samples were prepared by growing H99 to mid-logarithmic growth phase in three separate flasks of liquid YPD medium, with 150 rpm shaking. Approximately 1 × 10^9^ cells from each sample were pelleted, resuspended in fresh YPD medium, and incubated at 30°C for 90 min with 150 rpm shaking. Cells were then pelleted, flash frozen on dry ice, and lyophilized for ∼18 hours. Total RNA was isolated using the Qiagen RNeasy Plant Mini Kit (Qiagen, Valencia, CA); on-column DNase digestion was performed to ensure elimination of contaminating genomic DNA. Total RNA quantity and quality were assessed using the Agilent 2100 Bioanalyzer. Purified total RNA was subsequently stored at -80°C.

Aliquots from each total RNA sample were treated with one of three different RNA enrichment methods: the RNase H method for selective depletion of rRNA (Morlan *et al*. 2012; Adiconis *et al*. 2013), the Ribo-Zero rRNA Removal Kit (Yeast) (Illumina, San Diego, CA), and the NEBNext® Poly(A) mRNA Magnetic Isolation Module (NEB #E7490) (New England Biolabs, Ipswich, MA). RNA-Seq libraries were prepared from these enriched samples and from unenriched control “gold standard” samples (i.e. “Unenriched”) using the NEBNext® Ultra™ II Directional RNA Library Prep with Sample Purification Beads (NEB #E7765) and NEBNext® Multiplex Oligos for Illumina® (Dual Index Primers Set 1) (NEB #E7600) (New England Biolabs, Ipswich, MA). Libraries were pooled and sequenced by the Duke Sequencing and Genomic Technologies Shared Resource on an Illumina NextSeq 500 using the High-Output Kit to produce 75-basepair single-end reads.

It should be noted that while all work was done with the same three total RNA samples, the enrichment, library preparation, and sequencing were done in two batches, approximately one year apart. In between, total RNA samples were stored at - 80°C. Ribo-Zero-treated, Poly(A)-treated, and Unenriched RNA control libraries were prepared and sequenced in the first batch. RNase H-treated, replicate Poly(A)-treated, and replicate Unenriched RNA control libraries were prepared and sequenced in the second batch. All Unenriched RNA samples were compared and shown to be highly correlated (Figures 2A, 3A, & 4A), demonstrating that batch effect and differences in total RNA storage times did not confound comparisons. It should also be noted that the Ribo-Zero-treated libraries, the Poly(A)-treated libraries, and the Unenriched libraries were prepared by multiple individuals, which may explain some of the sample variation within those groups, while the RNase H-treated libraries were all prepared by a single individual. The individual who prepared each library is noted in the library metadata deposited at GEO.

### RNase H depletion

The RNase H depletion method (Morlan *et al*. 2012; Adiconis *et al*. 2013) is described briefly here; a more detailed protocol is included in supplementary data (File S1). The hybridization reaction mixture consisted of 1 µg of total RNA, 1 µl of 5x Hybridization Buffer (1000 mM NaCl, 500 mM Tris-HCl, pH 7.5), 0.65 µl of 100 µM pooled targeting oligos (discussed below), and nuclease-free water to bring the reaction to 5 µl. Oligo hybridization was performed in a thermocycler with the following program: 2 minutes at 95°C, ramp from 95°C to 22°C at -0.1°C/s, 5 minutes at 22°C. After hybridization, samples were transferred to ice and RNase H (New England Biolabs, Ipswich, MA) was added: 2 µl RNase H (5 U/µl), 1 µl 10x RNase H Reaction Buffer, 2 µl nuclease-free water. RNase H digestion was performed at 37°C for 30 minutes. After RNase H digestion, samples were stored on ice while adding DNase I (New England Biolabs, Ipswich, MA): 4 µl DNase I (2 U/µl), 10 µl 10x DNase I Reaction Buffer, 76 µl nuclease-free water. DNase I digestion was performed at 37°C for 30 minutes. After DNase I digestion, samples were transferred to ice. rRNA depleted RNA was purified from the reaction mixture with the Zymo RNA Clean & Concentrator-5 kit (Zymo Research, Irvine, CA) according to manufacturer instructions and eluted in 12 µl nuclease-free water. Finally, 5 µl of the eluted RNA was input to the library prep using NEBNext® Ultra II Directional RNA Library Prep Kit for Illumina (New England Biolabs, Ipswich, MA).

### Analysis overview

All genomic analyses used genome build CNA3 of H99 *Cryptococcus neoformans* var. *grubii* (accession GCA_000149245.3). The genome sequence and annotation were downloaded from release 39 of the Ensembl Fungi database (Kersey *et al*. 2016). For mapping mitochondrial rRNA genes, the original GTF downloaded from Ensembl Fungi was used. For all subsequent analyses, a modified GTF was used which included the newly mapped mitochondrial rRNA genes.

Analysis was performed using scripts written in the R programming language, Bash, and publicly available software detailed below. Custom R scripts used the following R and Bioconductor packages: Biostrings, BSgenome, dplyr, foreach, fs, GenomicAlignments, GenomicFeatures, ggbio, ggplot2, ggpubr, gridExtra, here, knitr, magrittr, plyr, readr, rmarkdown, Rsamtools, rstatix, rtracklayer, R.utils, stringr, tibble, tidyr, tools, utils.

### Mapping of mitochondrial rRNA genes

Coverage depth was plotted for all reads mapped to the mitochondrial chromosome for data generated from the first batch of Unenriched libraries. Visual inspection of these plots clearly indicated two regions with coverage depth several orders of magnitude higher than the rest of the chromosome. These regions do not overlap with any annotated feature in the mitochondrial chromosome. We determined the boundaries of these regions, extracted the sequences of the putative rRNA genes, and confirmed by BLASTn (Altschul *et al*. 1990) that these regions were homologous to known fungal mitochondrial small (positions 16948-18316) and large (positions 6710-9326) subunit rRNA genes (Figure S1). A modified version of the *C. neoformans* genome annotation supplemented with our mitochondrial rRNA gene annotations is included (File S2).

### Design of rRNA targeting oligonucleotides

Short DNA oligos were designed to target all nuclear rRNA genes (CNAG_10500, CNAG_10501, CNAG_10502, CNAG_10503) and the newly annotated 15S and 21S rRNA mitochondrial genes. In order to guide degradation of all rRNA by RNase H, the DNA oligos must be complementary to the rRNA and completely tile the rRNA. For simplicity and cost minimization, the design goal for rRNA targeting oligos was for them to be 50 nucleotides in length with no gaps between adjacent oligos. For genes with lengths that were not multiples of 50 nucleotides, single nucleotide gaps were introduced between oligos to allow for end-to-end coverage. Two, 55 nucleotide oligos were used to tile CNAG_10503, which is 111bp long.

Oligos were validated by mapping them to the H99 genome and confirming that they tiled as expected and mapped to the antisense strand. This validation process identified several partial duplications of the mitochondrial rRNA, putative nuclear mitochondrial DNA (numts) (Hazkani-Covo *et al*. 2010), and nuclear rRNA. These duplications were found in CNAG_04124, CNAG_06164, CNAG_07466, CNAG_12145, CNAG_12438, CNAG_13073, and in the region between CNAG_10503 and CNAG_03595. CNAG_13073 was excluded from analysis of rRNA depletion specificity because the rRNA duplication it contains is in an exon and on the sense strand, meaning that reads originating from rRNA genes can be misassigned to CNAG_13073. The other duplications do not result in spurious counts because they are either not in an exon or inserted antisense relative to the “host” gene.

The code used to design oligos should be applicable to other genomes; it is located within the file generate_rnaseh_oligos.Rmd, which is available, as described below, with the rest of the software developed for this project. This Rmarkdown document generates a TSV file in the correct format for pasting into the ordering template supplied by Eurofins; we have included a copy of the TSV generated for this project as a supplementary file (File S3). The 179 oligos were ordered from Eurofins Genomics LLC at a 10 nmol synthesis scale, with salt-free purification, resuspended to 100 µM, and shipped on dry ice. Upon receipt, all oligos were thawed, pooled, aliquoted, and stored at -80°C. Total cost for oligos (not including shipping) was less than $1000. This provided over 21 mL of pooled oligos (179 oligos at 120 µL per oligo), enough for over 33,000 reactions. Therefore, while the upfront cost of oligos is substantial, the per reaction cost is about $0.03.

### Bioinformatics and statistical data analyses

Basic assessments of sequence data quality were performed using FastQC (Andrews 2010) and MultiQC (Ewels *et al*. 2016). Raw sequencing reads were trimmed and filtered using fastq-mcf (EA-Utils version 1.04.807) (Aronesty 2011) and adapter sequences were extracted from the manufacturer-provided “Sample Sheet NextSeq E7600” template for the NEBNext® Multiplex Oligos for Illumina® (Dual Index Primers Set 1) (https://www.neb.com/tools-and-resources/usage-guidelines/sample-sheet-nextseq-e7600, accessed 11/10/2020). Reads were then mapped to the genome and read counts were generated using STAR (version 2.5.4b) (Dobin *et al*. 2013). For quantification of reads mapped to genes, we use the fourth column (“counts for the 2nd read strand aligned with RNA”) of the STAR ReadsPerGene.out.tab because the NEBNext® Ultra™ II Directional RNA Library Prep uses the dUTP method for strand-specific library preparation. All sequencing was done on an Illumina NextSeq 500, which has a flow cell with four lanes that are fluidically-linked (i.e., one pool is simultaneously loaded onto all four lanes). While we expect there to be some lane effects, we expect these to be less than fluidically-independent lanes. Because of this, and for simplicity, reads were combined across all four lanes for analysis of depletion efficiency and specificity.

### Analysis of rRNA depletion efficiency

To calculate the percentage of rRNA reads per library, Rsamtools (version 2.2.2) was used to extract reads from the STAR generated BAM files and determine the number of reads mapped to the rRNA genes. We did not use rRNA counts generated by STAR because STAR excludes multimapping reads from per gene read counts. As discussed above, several rRNA genes are partially duplicated elsewhere in the genome. Because STAR excludes multimapping reads from gene counts, it undercounts reads mapping to the rRNA genes that are partially duplicated. We confirmed the source of reads that mapped to rRNA duplicated regions by evaluating context: the count level of these reads corresponded to the level of expression of the rRNA genes from which the duplications seem to have arisen and not the level of expression of the genes (or genomic region in the case of the partial duplication of CNAG_10500) that seem to be the “acceptor sites” of these duplications. The percentage of total reads that mapped to rRNA genes was then calculated.

### Enrichment correlation analyses

Per gene read counts were generated by STAR as described above. Read counts for each library were combined across all four lanes using DESeq2::collapseReplicates, each library’s counts were normalized by its size factor, then an average count per gene was calculated for each enrichment method across all replicates. The mean normalized count of the Unenriched replicate libraries was considered the “gold standard” count for each gene. Specificity of each enrichment method was determined by calculating the Pearson correlation of the mean normalized count for each enrichment method with the Unenriched gold standard. Variation among the Unenriched libraries was quantified by cross-correlation: Pearson correlation was calculated for each Unenriched replicate library with the normalized mean of the other five Unenriched replicate libraries. Scatterplots were generated to visualize the correlation of replicate enriched libraries with the Unenriched gold standard. While calculation of mean normalized counts used all replicates for each method, scatterplots are only shown for one technical replicate of each RNA sample for each enrichment method. In addition to analyses across all genes, calculation of Pearson correlation and generation of scatterplots was repeated for subsets of genes, as annotated for “gene_biotype”: protein-coding genes, ncRNA, and tRNA, according to each gene’s annotation. rRNA genes and CNAG_13073 were excluded from all correlation analyses and scatterplots.

To determine specifically which genes seem to be “lost” by the Poly(A) isolation method, we identified genes with counts at least eight-fold lower in the Poly(A)-treated libraries than in the Unenriched libraries, after first excluding genes with very low expression in the Unenriched libraries (genes with less than 50 total read counts across all Unenriched libraries). These thresholds were chosen to identify obvious outliers in the Poly(A)-treated libraries, and were confirmed by visual inspection of the identified genes (Figure S6). We selected these thresholds to be more conservative than thresholds commonly used to identify genes with biologically relevant differences in expression (total reads of at least 10 and fold change of at least 2).

### LncRNA analysis

To identify novel lncRNA, we applied LncPipe (Zhao *et al*. 2018) to the data generated from the RNase H and Ribo-Zero enriched libraries and the Unenriched RNA libraries; Poly(A) enriched libraries were not included because they were expected to contain few, if any, reads derived from lncRNA. The published version of LncPipe only appears to work with data generated from human samples, so we forked the LncPipe repository and modified it to enable analysis of *C. neoformans* data. Details of the forked repository are provided below.

We developed an Rmarkdown document to perform all necessary pre-processing for running LncPipe (Zhao *et al*. 2018). This pre-processing involved automated reformatting of the input GTF file, preparing a subset of the GTF containing only protein-coding genes and another subset containing only non-protein-coding genes, generating a *C. neoformans* specific model for CPAT (one component of LncPipe), and generating a Bash script which itself runs LncPipe. LncPipe itself was run in Singularity with the bioinformatist/lncpipe Docker image built by the LncPipe developers (https://hub.docker.com/layers/bioinformatist/lncpipe/latest/images/sha256-9d97261556d0a3b243d4aa3eccf4d65e458037e31d9abb959f84b6fe54bb99a2?context=explore). Within LncPipe, STAR was used for mapping reads and the file step, LncPipeReporter, was not run.

### Data and reagent availability

The RNA-Seq data analyzed in this publication have been deposited in NCBI’s Gene Expression Omnibus (GEO) (Edgar *et al*. 2002) and will be accessible through GEO Series accession number GSE160397 (https://www.ncbi.nlm.nih.gov/geo/query/acc.cgi?acc=GSE160397). The custom programs developed for processing and analyzing the RNA-Seq data are available in a GitHub repository (https://github.com/granek/rna_enrichment) and the version of the LncPipe pipeline that we modified to run on the H99 genome is available in a GitHub repository (https://github.com/granek/LncPipe) that was forked from the original. For purposes of reproducibility, all analyses were run within Singularity containers (v 3.5.2). All lncRNA discovery was performed using the bioinformatist/lncpipe Docker image (run within Singularity) provided by the LncPipe developers (https://hub.docker.com/layers/bioinformatist/lncpipe/latest/images/sha256-9d97261556d0a3b243d4aa3eccf4d65e458037e31d9abb959f84b6fe54bb99a2?context=explore). All other analyses were performed using a Singularity image which we built and is publicly available (library://granek/published/rna_enrichment). These resources include all programs, support files, and instructions for automatically replicating all analyses presented here using the data available from GEO.

Figure S1 contains a depth of coverage plot of the mitochondrial rRNA genes. Figures S2, S3, and S4 display scatterplot visualizations of rRNA depletion specificity summarized in Figures 2, 3, and 4, respectively. Figure S5 displays the rRNA depletion efficiency for ncRNA genes, excluding CNAG_12993. Figure S6 displays a scatterplot visualization of the genes that are underrepresented by the Poly(A) isolation method and Table S1 provides details of these genes. File S1 contains the RNase H depletion protocol. File S2 contains the Ensembl GTF with newly annotated mitochondrial rRNA. File S3 contains DNA oligonucleotide sequences used in the RNase H depletion method. All supplementary information has been deposited in figshare.

## RESULTS

### RNase H depletion is most efficient in removing rRNA

We focus the majority of our analyses on the RNase H depletion and Poly(A) isolation methods, because, of the three RNA enrichment methods assessed here, they are the two that are still available for use. To provide some context to our RNase H depletion method results, we also include analyses on the Ribo-Zero depletion method, which was frequently used in fungal RNA-Seq experiments before its discontinuation.

As an initial assessment, we evaluated the efficiency with which each enrichment method removed rRNA. To do so, we quantified the percentage of total reads that mapped to rRNA genes for each method and compared these percentages to those of Unenriched RNA control libraries generated by sequencing identical RNA samples without any enrichment. As expected, the vast majority (∼90-92%) of reads in the Unenriched RNA control libraries map to rRNA genes (Figure 1). Both the RNase H-treated libraries (∼1.5-2.5%) and the Poly(A)-treated libraries (∼3-5%) display a significant reduction in the percentage of reads mapping to rRNA genes (Figure 1). The Ribo-Zero depletion method was previously demonstrated to be efficient in depleting fungal rRNA and was used successfully in RNA-Seq applications for various fungi (Illumina; Trevijano-Contador *et al*. 2018; Liu *et al*. 2020). We similarly evaluated the Ribo-Zero depletion method and observed that the number of mapped rRNA reads is significantly higher in the Ribo-Zero-treated libraries (∼21-85%) than in the RNase H-treated and Poly(A)-treated libraries (Figure 1). Overall, both the RNase H depletion and Poly(A) isolation methods demonstrate robust efficiency in removing fungal rRNA, with the RNase H depletion method modestly outperforming the commonly-used Poly(A) isolation method.

**Figure 1.**
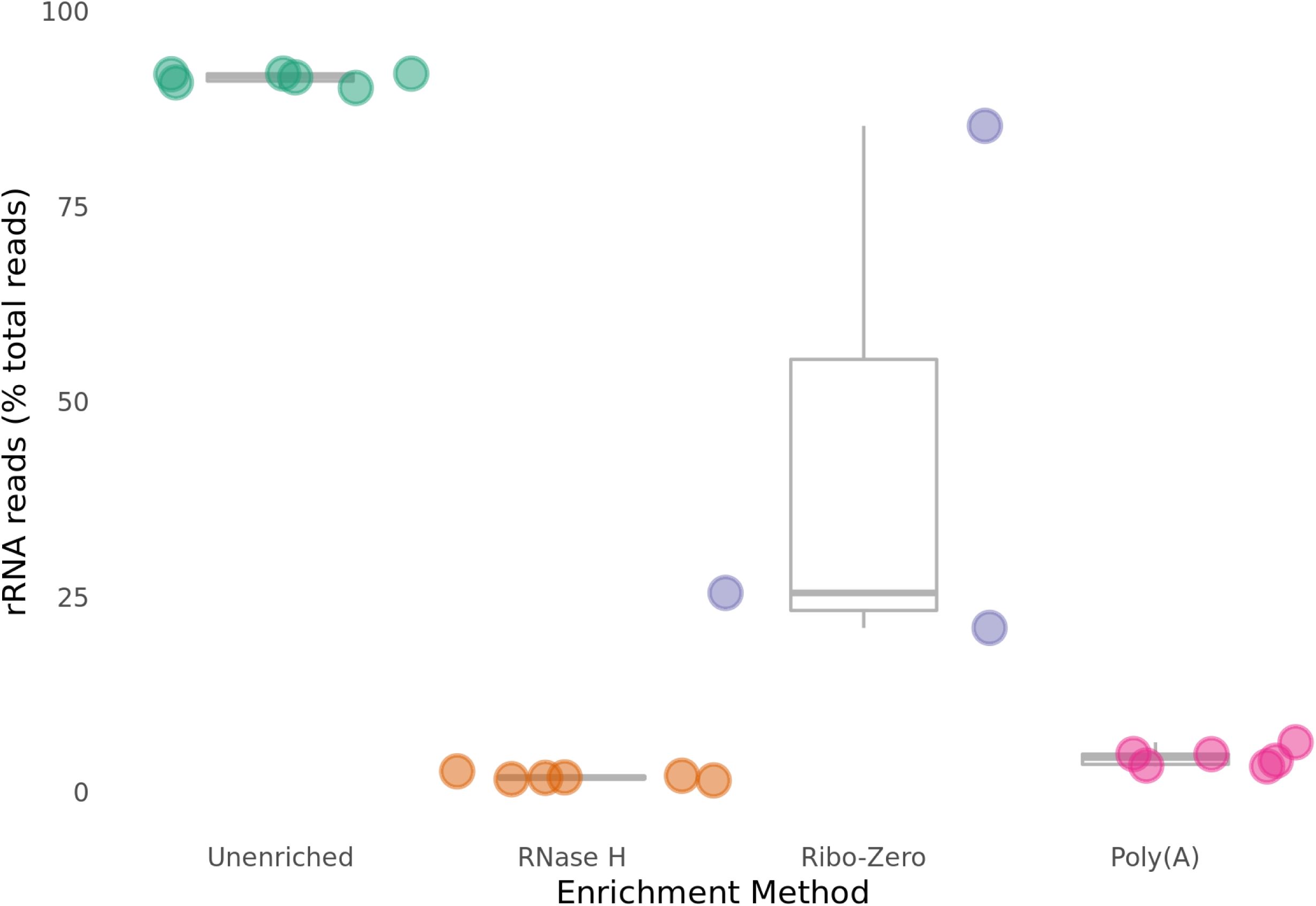
rRNA depletion efficiency: The percentage of rRNA reads in each library is graphed. The RNase H depletion method has the most efficient depletion (lowest percentage of rRNA reads), with the Poly(A) isolation method a close second, and the Ribo-Zero depletion method a distant third. Unenriched libraries show that rRNA makes up most of the RNA in *C. neoformans*.

### RNase H depletion more accurately preserves true levels of non-rRNA than Poly(A) isolation

To compare the specificity of the three RNA enrichment methods, we determined the correlation between read counts in the enriched libraries to read counts in the Unenriched RNA control libraries generated from the same samples. To do so, we first calculated the correlation coefficient between normalized reads mapped to all non-rRNA genes from all Unenriched RNA samples, in order to determine the maximum achievable correlation between libraries. As expected, we observed that the Unenriched RNA samples are highly correlated (R = 0.983-0.997), demonstrating reproducibility between samples and batches (Figure 2A).

**Figure 2.**
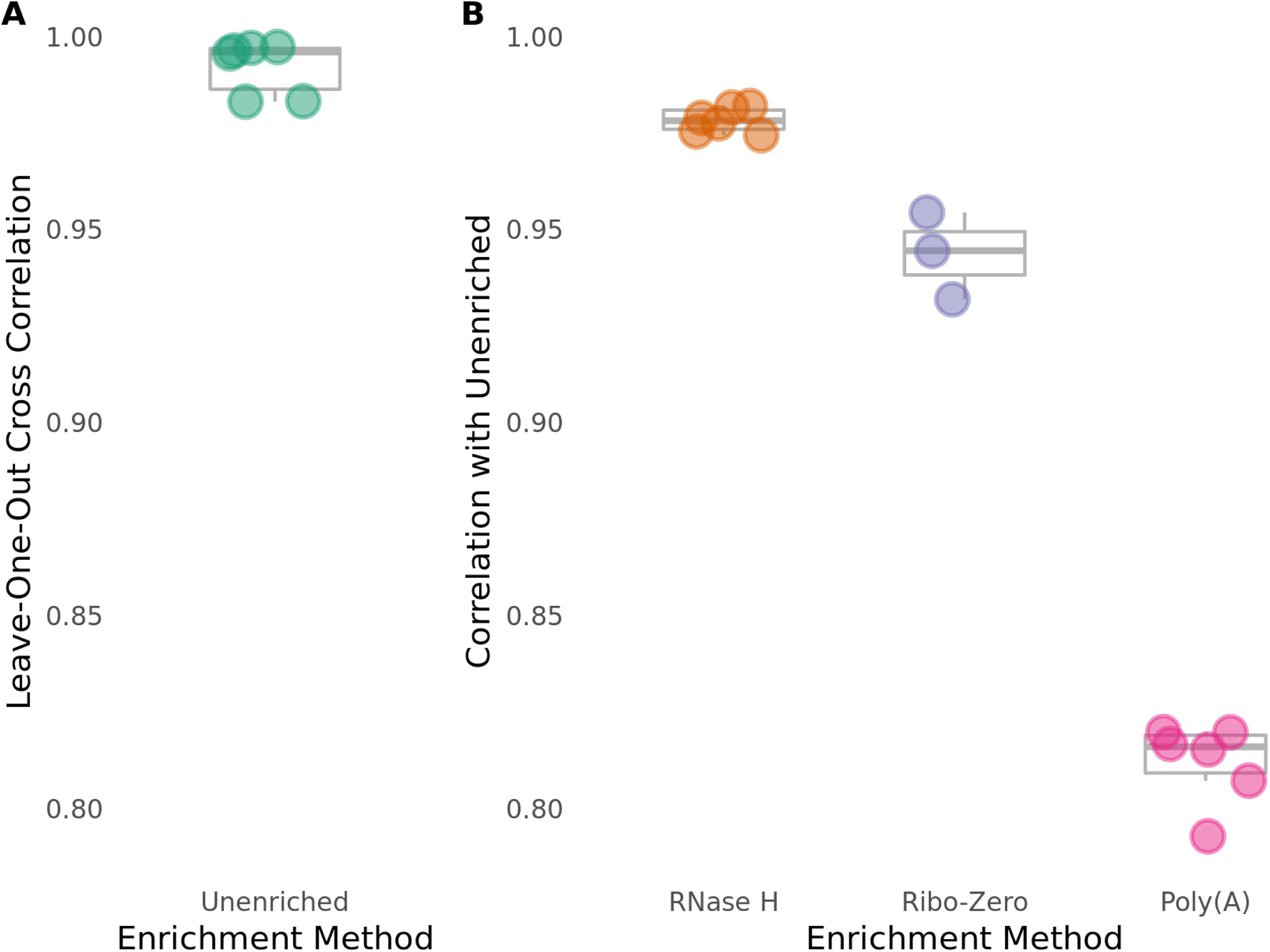
Specificity of rRNA depletion for all genes: Pearson correlations were calculated for normalized read counts of all annotated genes in the *C. neoformans* genome, excluding rRNA genes and genes containing coding-strand rRNA duplications. A. Unenriched libraries have high internal consistency as determined by leave-one-out cross correlation of each Unenriched library with the mean of other Unenriched libraries. B. The RNase H depletion method has the best overall rRNA depletion specificity, as determined by Pearson correlation of read counts for all genes with the Unenriched libraries. Pearson correlation coefficient (R) was calculated between each enriched library and the gold standard, the gene-wise average of counts across all Unenriched library replicates.

We compared the abilities of the RNase H depletion method and the Poly(A) isolation method to preserve all non-rRNA following rRNA depletion. To do so, we calculated the correlation coefficient between normalized reads mapped to all non-rRNA genes in RNase H-treated/Poly(A)-treated libraries and normalized reads mapped to all non-rRNA genes from the Unenriched RNA control libraries. We found that the RNase H-treated libraries (R = 0.974-0.982) display a much better correlation with the Unenriched RNA control libraries than the Poly(A)-treated libraries (R = 0.793-0.820) for all reads mapping to non-rRNA genes (Figures 2B & S2). We similarly assessed the Ribo-Zero depletion method for preservation of all non-rRNA. Ribo-Zero-treated libraries (R = 0.932-0.954) display a much better correlation with the Unenriched libraries than the Poly(A)-treated libraries, but a slightly weaker correlation than the RNase H-treated libraries for reads mapping to all non-rRNA genes (Figure 2B & S2). This observation suggests that the RNase H depletion method may be more specific than the Poly(A) isolation and the Ribo-Zero depletion methods, in that it maintains non-rRNA levels observed in the Unenriched RNA control libraries.

### RNase H depletion more accurately preserves true levels of protein-coding RNA than Poly(A) isolation

We next assessed the ability of each RNA enrichment method to retain protein-coding RNA specifically. To do so, we calculated the correlation coefficient between normalized reads mapped to protein-coding genes from all Unenriched RNA samples, in order to determine the maximum achievable correlation between libraries. As expected, we observed that the Unenriched RNA samples are highly correlated (R = 0.986-0.998), demonstrating reproducibility between samples and batches (Figure 3A).

We compared the ability of the RNase H depletion method and the Poly(A) isolation method to preserve protein-coding RNA following rRNA depletion. To do so, we calculated the correlation coefficient between normalized reads mapped to protein-coding genes in RNase H-treated/Poly(A)-treated libraries and normalized reads mapped to protein-coding genes from the Unenriched RNA control libraries. We found that the RNase H-treated libraries (R = 0.985-0.990) display a much better correlation with the Unenriched RNA control libraries than the Poly(A)-treated libraries (R = 0.810-0.838) for all reads mapping to protein-coding genes (Figures 3B & S3). We similarly assessed the Ribo-Zero depletion method for preservation of protein-coding RNA. Ribo-Zero-treated libraries (R = 0.935-0.962) display a better correlation with the Unenriched libraries than the Poly(A)-treated libraries, but a slightly weaker correlation than the RNase H-treated libraries for reads mapping to protein-coding genes (Figure 3B & S3). This observation demonstrates that the RNase H depletion method is more specific than both the Poly(A) isolation and Ribo-Zero depletion methods in preserving protein-coding RNA levels observed in the Unenriched RNA control libraries.

**Figure 3.**
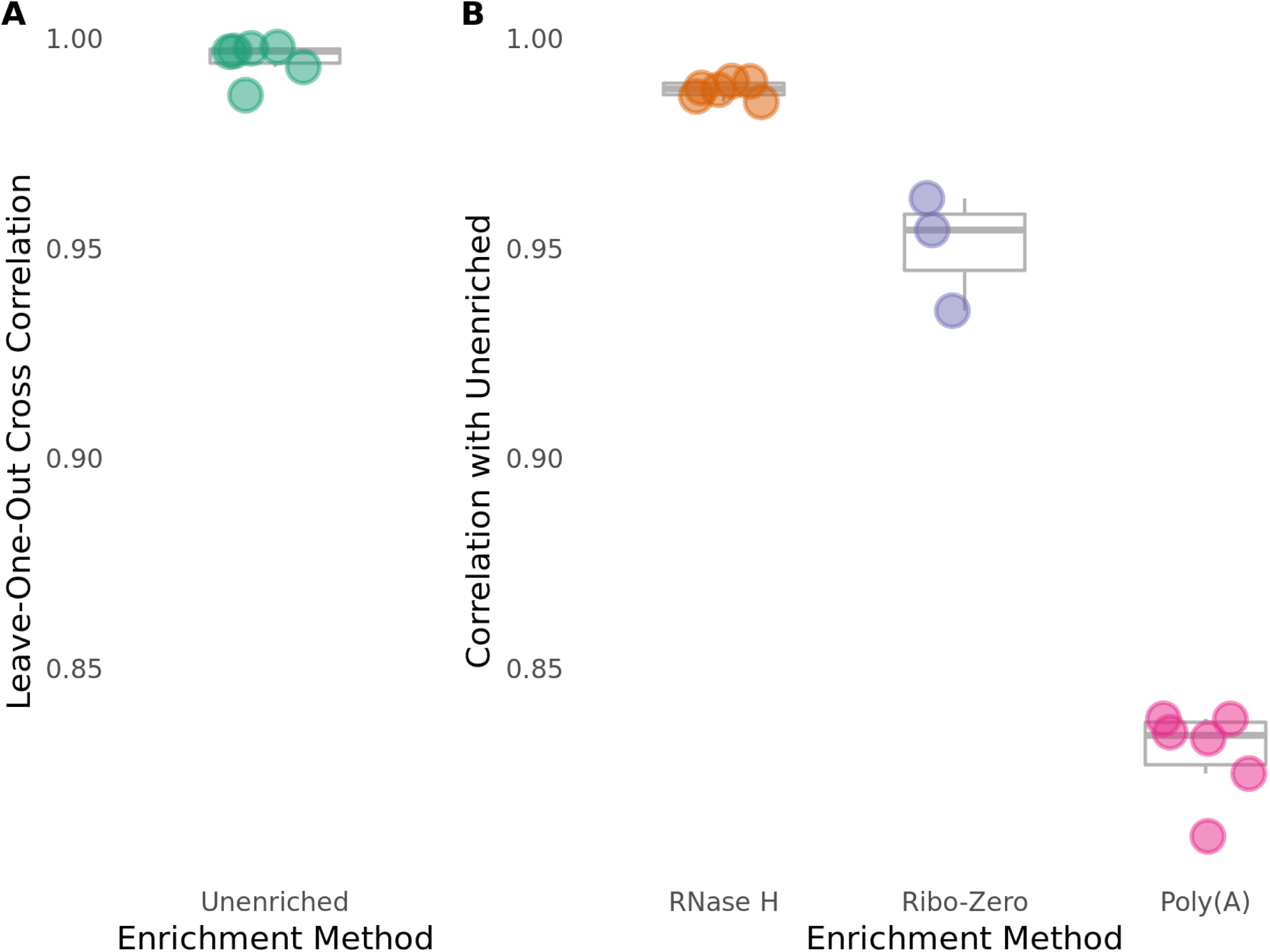
Specificity of rRNA depletion for protein-coding genes: Pearson correlations were calculated in the same way as Figure 2, but only for protein-coding genes, excluding genes containing coding-strand rRNA duplications. A. Unenriched libraries have high internal consistency for protein-coding genes. B. The RNase H depletion method has the best rRNA depletion specificity for protein-coding genes.

### RNase H depletion more accurately preserves true levels of ncRNA than Poly(A) isolation, but slightly less accurately than Ribo-Zero depletion

We next assessed the ability of each RNA enrichment method to retain ncRNA specifically. To do so, we calculated the correlation coefficient between normalized reads mapped to ncRNA genes from all Unenriched RNA samples, in order to determine the maximum achievable correlation between libraries. Again, we observed that the Unenriched RNA samples are highly correlated (R = 0.835-0.990), demonstrating reproducibility between samples and batches (Figure 4A).

**Figure 4.**
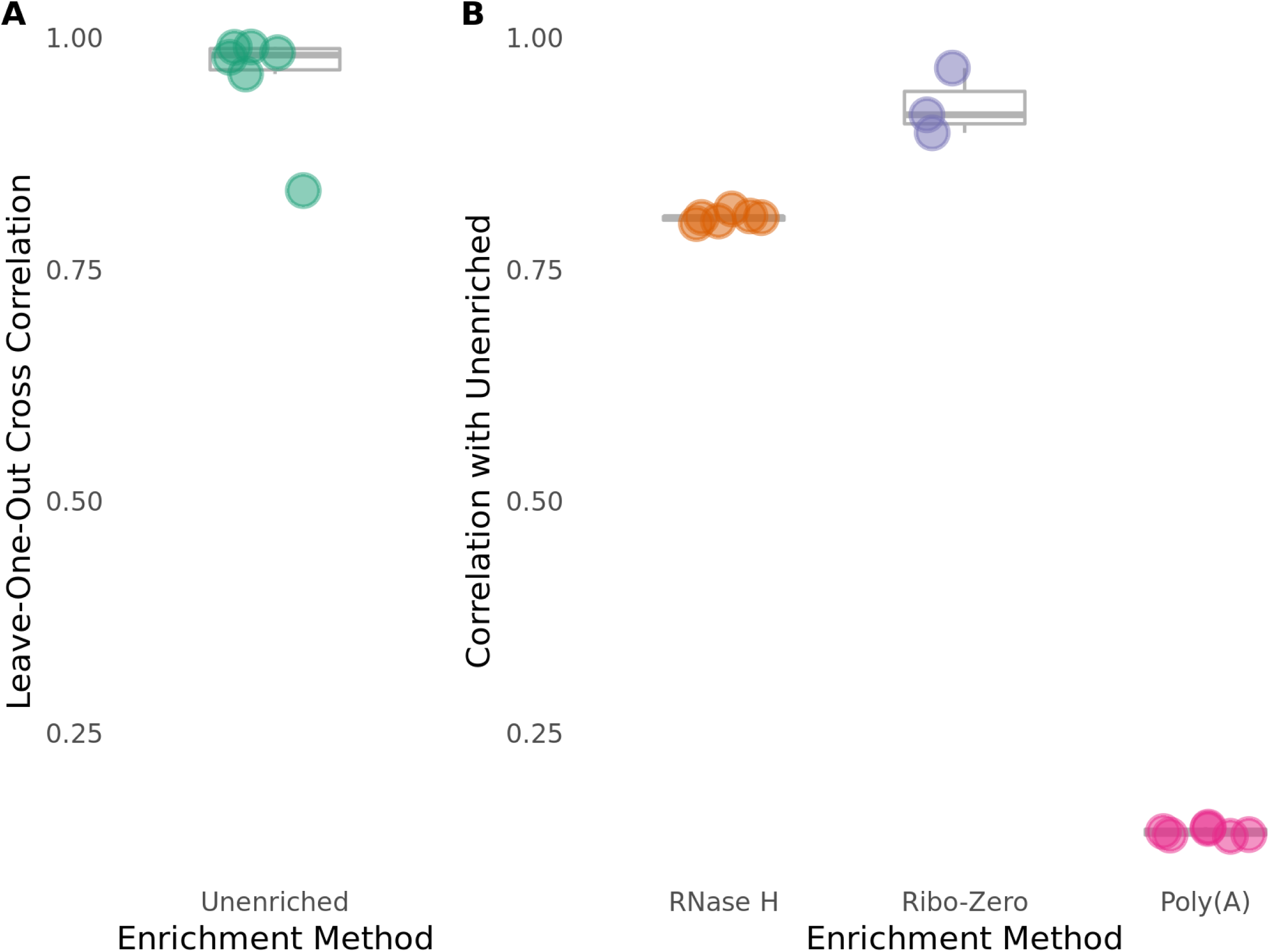
Specificity of rRNA depletion for ncRNA genes: Pearson correlations were calculated in the same way as Figure 2, but only for ncRNA genes, excluding rRNA genes. A. Unenriched libraries have high internal consistency for ncRNA genes. B. The Ribo-Zero depletion method has the best rRNA depletion specificity for ncRNA genes.

We compared the ability of the RNase H depletion method and the Poly(A) isolation method to preserve ncRNA following rRNA depletion. We calculated the correlation coefficient between normalized reads mapped to ncRNA genes in RNase H-treated/Poly(A)-treated libraries and normalized reads mapped to ncRNA genes from the Unenriched RNA control libraries. As expected, we found that the RNase H-treated libraries (R = 0.799-0.815) display a much better correlation with the Unenriched RNA control libraries than the Poly(A)-treated libraries (R = 0.139-0.149) for all reads mapping to ncRNA genes (Figures 4B & S4). This result was expected because the Poly(A) isolation method specifically enriches RNA with polyadenylation and excludes all other non-polyadenylated RNA, including ncRNA and tRNA. One key advantage of methods that specifically remove rRNA, such as the RNase H depletion and the Ribo-Zero depletion methods, is that they “ignore” all RNA that is not specifically targeted for removal. As a result, these non-polyadenylated RNA species should maintain similar levels as the input Unenriched RNA.

As a better assessment of the ability of the RNase H depletion method to preserve ncRNA, we compared it to the Ribo-Zero depletion method. Ribo-Zero-treated libraries (R = 0.897-0.967) display a slightly better correlation with the Unenriched libraries than the RNase H-treated libraries for reads mapping to ncRNA genes (Figure 4B & S4). The higher correlation of Ribo-Zero-treated libraries with Unenriched libraries seems to be driven by CNAG_12993, the ncRNA gene with the highest counts in the Unenriched libraries, but much lower counts in the RNase H libraries. When CNAG_12993 is removed from analysis, the RNase H and Ribo-Zero depletion methods perform similarly (Figure S5; RNase H R = 0.936-0.952, Ribo-Zero R = 0.892-0.959). There is no clear explanation for the poor performance of the RNase H depletion method with CNAG_12993.

We also explored the ability of each enrichment method to preserve tRNA. Fewer than 10 reads mapped to each tRNA gene in all libraries, likely due to size selection in the library preparation, precluding any meaningful analysis (data not shown).

### LncPipe pipeline identifies novel lncRNA in the *C. neoformans* transcriptome

Because our correlation analyses demonstrated that both the RNase H depletion and the Ribo-Zero depletion methods preserve ncRNA, we took advantage of this rich RNA-Seq dataset to search for novel lncRNAs. We used an existing pipeline LncPipe (Zhao *et al*. 2018) that was developed for a subset of model organisms, and modified it for application to *C. neoformans*. We applied this modified LncPipe pipeline to identify novel lncRNA within our RNase H depletion, Ribo-Zero depletion, and Unenriched RNA datasets. Our lncRNA discovery analysis identified 11 novel lncRNA within the *C. neoformans* transcriptome (Table 1).

**Table 1.** LncPipe identification of novel *C. neoformans* lncRNA: Putative lncRNAs were discovered by analysis of RNase H-treated, Ribo-Zero-treated, and Unenriched RNA libraries. The name (assigned by LncPipe), chromosomal location, exon number, exonic length, and transcripts per million (TPM) across samples are shown for all 11 novel lncRNA identified.

| Name | Chromosome | Start | End | # Exons | Total Exonic Length | Mean TPM | Median TPM |
| --- | --- | --- | --- | --- | --- | --- | --- |
| LINC-CNAG_07358-1 | 1 | 996421 | 997387 | 2 | 863 | 6.689427833 | 6.2667 |
| LINC-CNAG_07633-1 | 6 | 499352 | 499840 | 3 | 350 | 2.844233167 | 0 |
| LINC-CNAG_07649-1 | 6 | 1351673 | 1352718 | 3 | 913 | 5.5161474 | 5.799715 |
| LINC-CNAG_07769-5 | 9 | 828268 | 829327 | 3 | 919 | 9.2005976 | 10.76355 |
| LINC-CNAG_07769-4 | 9 | 831951 | 833064 | 4 | 2019 | 5.333811583 | 5.026716 |
| LINC-CNAG_07769-1 | 9 | 838389 | 840007 | 10 | 2730 | 4.443440783 | 3.846055 |
| LINC-CNAG_04857-1 | 10 | 199988 | 203380 | 43 | 6849 | 12.5665295 | 12.729905 |
| LINC-CNAG_04857-2 | 10 | 203693 | 205767 | 2 | 1983 | 1.903616517 | 1.67792 |
| LINC-CNAG_01945-1 | 11 | 1333989 | 1334590 | 2 | 540 | 3.47618165 | 1.745465 |
| LINC-CNAG_06521-2 | 13 | 743402 | 744570 | 4 | 1003 | 5.924291983 | 5.627535 |
| LINC-CNAG_07042-1 | 13 | 750280 | 751009 | 3 | 609 | 5.950332833 | 4.02392 |

## DISCUSSION

RNA enrichment is essential for cost-effectively generating data from an RNA-Seq experiment. We have demonstrated here that, in *C. neoformans* cells grown in rich media, rRNA constitutes more than 90% of the total RNA; even higher percentages of rRNA have been observed in other species (Giannoukos *et al*. 2012). RNA-Seq experiments are typically aimed at quantifying protein-coding RNA, and increasingly also ncRNA. Efficient reduction of rRNA allows one to generate the desired sequencing depth of the RNA species of interest with one-tenth of the sequencing reads that would be required to generate the same depth from total, unenriched RNA.

To be effective, RNA enrichment methods must be efficient and specific. An efficient RNA enrichment method removes as much rRNA as possible. A specific RNA enrichment method does not affect other RNA species in the sample. We compared the rRNA removal efficiency of three commonly-used methods in *C. neoformans* samples. Application of the RNase H depletion method in *Cryptococcus* has never been reported to our knowledge. The Poly(A) isolation method (Bloom *et al*. 2019; Brown *et al*. 2020) and the now discontinued Ribo-Zero depletion method have both been used in RNA-Seq applications with *Cryptococcus* samples in the past (Illumina; Trevijano-Contador *et al*. 2018; Liu *et al*. 2020). We find that both the untested RNase H depletion method, as well as the frequently-used Poly(A) selection method, are very efficient in removing fungal rRNA. Surprisingly, the Ribo-Zero depletion method showed poor efficiency in *C. neoformans*, despite previous work showing efficient removal of various bacterial rRNA (Giannoukos *et al*. 2012). While the Ribo-Zero manufacturer predicted that the Ribo-Zero Yeast kit would work for *C. neoformans*, the probes were designed to target *S. cerevisiae*, which may explain the poor performance observed here. Of the three methods tested, the RNase H depletion method is the most efficient in removing fungal rRNA.

Following the removal of rRNA, the majority of remaining RNA is typically protein-coding. Ideally, the removal of rRNA should not have any effect on protein-coding RNA. In reality, there is no known method that can reduce rRNA without having some effect on non-target RNA, including protein-coding RNA. The best available methods efficiently remove rRNA and are highly specific, having minimal side-effects on non-target RNA. When assessing the ability of these RNA enrichment methods to preserve protein-coding RNA, we observe that the RNase H depletion method is substantially more specific than the Poly(A) isolation method and somewhat more specific than the Ribo-Zero depletion method.

We evaluated the specificity of each RNA enrichment method in preserving ncRNA. The Poly(A) isolation method performs very poorly in preserving ncRNA; this is as expected since it depends on 3’ polyadenylation, which is absent from ncRNA. The RNase H depletion and the Ribo-Zero depletion methods both perform well in preserving ncRNA, with the Ribo-Zero depletion method being slightly more specific than the RNase H depletion method.

To determine a possible mechanism impacting specificity of the Poly(A) isolation method, we identified the genes that were significantly underrepresented in the Poly(A)-treated libraries compared to the Unenriched libraries (Figure S6). A total of 41 genes were identified as significantly underrepresented in the Poly(A)-treated libraries; as expected, 24 of these genes are non-coding genes (Table S1). In analyzing the remaining 17 protein-coding genes that were underrepresented in the Poly(A)-treated libraries, we observed a pattern. The vast majority of these genes (12 of 17) are located on the mitochondrial chromosome (Table S1). This observation is supported by previous work that has demonstrated that mitochondrial transcripts in fungi, including *Cryptococcus*, lack polyadenylation (Toffaletti *et al*. 2003; Chang and Tong 2012). From this analysis, it appears that the Poly(A) isolation method is largely specific for most protein-coding genes except for mitochondrial genes. We have not identified any characteristic that explains the five underrepresented protein coding genes that reside in the nuclear genome. These findings indicate that the Poly(A) isolation method is not well suited for RNA-Seq experiments investigating ncRNA genes, genes on the mitochondrial chromosome, or the five nuclear protein-coding genes that we found to be underrepresented in the Poly(A)-treated libraries.

Interest in ncRNA has recently expanded in the fungal genetics field. The majority of work has focused on ncRNAs in model systems (such as *S. cerevisiae, Neurospora crassa*, and *Aspergillus flavus*), in which lncRNAs and natural antisense transcripts (NATs) have been implicated in stress responses and development (Smith *et al*. 2008; Gelfand *et al*. 2011; Ding *et al*. 2012; Xue *et al*. 2014). Little work has explored ncRNAs in pathogenic fungi. For example, RNA interference (RNAi) is known to regulate transposon activity in *C. neoformans* (Wang *et al*. 2010; Janbon *et al*. 2010; Yadav *et al*. 2018). The first lncRNA in *Cryptococcus*, *RZE1*, was recently functionally characterized. *RZE1* is required for *Cryptococcus* yeast-to-hyphal transition and virulence through its regulation of the transcription factor *ZNF2* (Chacko *et al*. 2015). Additionally, siRNAs, miRNAs, and lncRNAs are known to be secreted, albeit for an unknown purpose, by *C. deneoformans* (Liu *et al*. 2020). We have identified 11 putative lncRNAs in *C. neoformans* by mining our dataset by modifying LncPipe (Zhao *et al*. 2018) to run on genomic data from non-model organisms. Although many of these lncRNAs were not highly expressed in the evaluated condition, they may be interesting candidates to pursue for novel biological activity in conditions relevant to fungal pathogenesis, because many fungal ncRNAs are induced in response to stressful stimuli. Furthermore, the *C. neoformans* genome may contain undiscovered lncRNAs that are not expressed in the rich growth conditions used here.

In conclusion, of the three RNA enrichment methods compared here, we consider the Poly(A) isolation method to have poor overall effectiveness. It does efficiently reduce rRNA reads (although not as efficiently as the RNase H depletion method), but it results in a biased enrichment of protein-coding transcripts. The RNase H depletion and the Ribo-Zero depletion methods both display strengths and weaknesses. The RNase H depletion method performs better in efficiency of rRNA reduction and specificity for protein-coding transcripts, while the Ribo-Zero depletion method performs moderately better in specificity for ncRNA. While this work was being conducted, the Ribo-Zero product line was discontinued. It has since been replaced with the Illumina Ribo-Zero Plus rRNA Depletion Kit, which only targets human, mouse, rat, and bacterial rRNA. As a result, we conclude that the RNase H method may be the best option for RNA-Seq analysis of *C. neoformans*, as well as many other non-model organisms. While the RNase H depletion method has a substantial upfront cost to purchase DNA oligonucleotides (approximately $1000), we estimate that for this method our total cost per sample was less than $6.50 (more than half of this total was the final cleanup with the Zymo RNA Clean & Concentrator-5 kit).

## ACKNOWLEDGEMENTS

We thank the Duke University School of Medicine for the use of the Microbiome Shared Resource, which provided RNA quality assessments. We also thank the Duke University School of Medicine for the use of the Sequencing and Genomic Technologies Shared Resource, which provided all sequencing services used for this project. This work was made possible by funding from the National Institute of Biomedical Imaging and Bioengineering (grant #5R25EB023928) received by KO and CC, and R01 grant AI074677 to JAA and JAG.

## SUPPLEMENTARY MATERIAL

**Figure S1. Visualization of mitochondrial rRNA genes in *C. neoformans* H99 strain:** The depth of coverage plot of the whole H99 mitochondrial chromosome is shown. The *C. neoformans* large and small mitochondrial rRNA genes are clearly identified as the two regions of the mitochondrial genome with depth of coverage that is much higher than any other part of the chromosome.

**Figure S2. Scatterplot visualization of rRNA depletion specificity for all genes:** In the RNase H- and Ribo-Zero-treated libraries, most genes have counts that are highly correlated with the Unenriched libraries, whereas Poly(A)-treated libraries have low counts for a number of expressed genes. For each biological replicate (subplot columns labeled “A”, “B”, and “C”), per-gene normalized read counts for each enrichment method are plotted as a function of the gold standard (normalized read counts averaged across the Unenriched libraries). All annotated genes in the *C. neoformans* genome are plotted, excluding rRNA genes and genes containing coding-strand rRNA duplications. For libraries with technical replicates (RNase H and Ribo-Zero), only one of the replicates is shown.

**Figure S3. Scatterplot visualization of rRNA depletion specificity for protein-coding genes:** In the RNase H- and Ribo-Zero-treated libraries, most protein-coding genes have counts that are highly correlated with the Unenriched libraries, whereas Poly(A)-treated libraries have low counts for a number of protein-coding genes. This plot is the same as Figure S2, but only shows protein-coding genes, excluding genes containing coding-strand rRNA duplications.

**Figure S4. Scatterplot visualization of rRNA depletion specificity for ncRNA genes:** In the RNase H- and Ribo-Zero-treated libraries, most ncRNA genes have counts that are well-correlated with the Unenriched libraries, whereas Poly(A)-treated libraries have low counts for almost all ncRNA genes. This plot is the same as Figure S2, but only shows ncRNA genes, excluding rRNA.

**Figure S5. Specificity of rRNA depletion for ncRNA genes, excluding CNAG_12993:** When outlier ncRNA gene CNAG_12993 is excluded, the RNase H depletion method has specificity for ncRNA genes that is as good as the Ribo-Zero depletion method. Pearson correlations were calculated in the same way as Figure 4, except CNAG_12993 was excluded from the analysis.

**Figure S6. Scatterplot visualization of all genes underrepresented by the Poly(A) isolation method:** This plot is the same as Figure S2, with the genes that are significantly underrepresented in the Poly(A)-treated libraries colored red.

**Table S1. Genes underrepresented by the Poly(A) isolation method:** Table details the genes that are significantly underrepresented in the Poly(A)-treated libraries. The gene name, chromosome, and gene type are included for each gene.

**File S1. RNase H depletion protocol:** The detailed protocol used to perform the RNase H depletion.

**File S2. Ensembl GTF with newly annotated mitochondrial rRNA:** A copy of the GTF genome annotation file for CNA3 of H99 *Cryptococcus neoformans* var. *grubii*, modified to include annotation for the mitochondrial rRNA genes.

**File S3. RNase H rRNA-targeting oligonucleotides:** DNA oligonucleotide sequences used in the RNase H depletion method to target H99 *C. neoformans* rRNA. This file was formatted to be directly pasted into the Eurofins order form.

